# Joint analysis of functional genomic data and genome-wide association studies of 18 human traits

**DOI:** 10.1101/000752

**Authors:** Joseph K. Pickrell

## Abstract

Annotations of gene structures and regulatory elements can inform genome-wide association studies (GWAS). However, choosing the relevant annotations for interpreting an association study of a given trait remains challenging. We describe a statistical model that uses association statistics computed across the genome to identify classes of genomic element that are enriched or depleted for loci that influence a trait. The model naturally incorporates multiple types of annotations. We applied the model to GWAS of 18 human traits, including red blood cell traits, platelet traits, glucose levels, lipid levels, height, BMI, and Crohn’s disease. For each trait, we evaluated the relevance of 450 different genomic annotations, including protein-coding genes, enhancers, and DNase-I hypersensitive sites in over a hundred tissues and cell lines. We show that the fraction of phenotype-associated SNPs that influence protein sequence ranges from around 2% (for platelet volume) up to around 20% (for LDL cholesterol); that repressed chromatin is significantly depleted for SNPs associated with several traits; and that cell type-specific DNase-I hypersensitive sites are enriched for SNPs associated with several traits (for example, the spleen in platelet volume). Finally, by re-weighting each GWAS using information from functional genomics, we increase the number of loci with high-confidence associations by around 5%.

## 1 Introduction

A fundamental goal of human genetics is to create a catalogue of the genetic polymorphisms that cause phenotypic variation in our species and to characterize the precise molecular mechanisms by which these polymorphisms exert their effects. An important tool in the modern human genetics toolkit is the genome-wide association study, in which hundreds of thousands or millions of single nucleotide polymorphisms (SNPs) are genotyped in large cohorts of individuals and each polymorphism tested for a statistical association with some trait of interest. In recent years, GWAS have identified thousands of genomic regions that show reproducible statistical associations with a wide array of phenotypes and diseases^1^.

In general, the loci identified in GWAS of multifactorial traits have small effect sizes and are located outside of protein-coding exons^2^. This latter fact has generated considerable interest in annotating other types of genomic elements apart from exons. For example, the ENCODE project has generated detailed maps of histone modifications and transcription factor binding in six human cell lines, partially motivated by the goal of interpreting GWAS signals that may act via a mechanism of gene regulation^3^. Methods for combining potentially rich sources of functional genomic data with GWAS could in principle lead to important biological insights. The development of such a method is the aim of this paper.

There are two lines of research that motivate our work on this problem. The first is what are often called “enrichment” analyses. In this type of analysis, the most strongly associated SNPs in a GWAS are examined to see if they fall disproportionately in specific types of genomic region. These studies have found, for example, that SNPs identified in GWAS are enriched in protein-coding exons, promoters, and untranslated regions (UTRs)^2;4^ and among those that influence gene expression^5;6^. Further, in some cases, SNPs associated with a trait are enriched in gene regulatory regions in specific cell types^7–18^ or near genes expressed in specific cell types^19;20^. However, the methods in these studies are generally not able to consider more than a single annotation at a time (with a few exceptions^21;22^). Further, they are not set up to answer a question that we find important: consider two independent SNPs with equivalent P-values of 1 × 10^−7^ in a GWAS for some trait (note that this P-value does not reach the standard threshold of 5 × 10^−8^ for “significance”), the first of which is a nonsynonymous SNP and the second of which falls far from any known gene. What is the probability that the first SNP is truly associated with the trait, and how does this compare to the probability for the second?

A potential answer to this question comes from the second line of research that motivates this work. In association studies where the phenotype being studied is gene expression (“eQTL” studies, for “expression quantitative trait locus”), statistical models have been developed to identify shared characteristics of SNPs that influence gene expression^23–25^. In a hierarchical modeling framework, the probability that a given SNP influences gene expression can then depend on these characteristics. The key fact that makes these models useful in the context of eQTL mapping is that there is a large number of unambiguous eQTLs in the genome on which a model can be trained. In the GWAS context, the number of loci unambiguously associated with a given trait has historically been very small; learning the shared properties of two or three loci is not a job well-suited to statistical modeling. However, large meta-analyses of GWAS now regularly identify tens to hundreds of independent loci that influence a trait (e.g.^26;27^). The merits of hierarchical modeling in this context^28–30^ are thus worth revisiting. Indeed, Carbonetto and Stephens^31^ have reported success in identifying loci involved in autoimmune diseases using a hierarchical model that incorporates information about groups of genes known to interact in a pathway.

In this paper we present a hierarchical model for jointly analyzing GWAS and genomic annotations. We applied this model to GWAS of 18 diseases and traits; for each trait, we learned the relevant types of genomic information from a set of 450 genome annotations.

## 2 Results

We assembled a set of 18 GWAS with publicly available summary statistics and a large number (at least around 20) of loci associated with the trait of interest. These included studies of red blood cell traits^15^, platelet traits^32^, Crohn’s disease^33^, BMI^34^, lipid levels^27^, height^26^, bone mineral density^35^ and fasting glucose levels^36^. We used ImpG^37^ to impute the summary statistics from each study for all common SNPs identified in European populations by the 1000 Genomes Project^38^. Overall we successfully imputed association statistics for around 80% of common SNPs (Supplementary Figure 1). We then assembled a set of genome annotations, paying specific attention to annotations available for many cell types since important regulatory elements may be active only in specific cell types. The main sources of genome annotations were 402 maps of DNase-I hypersensitivity in a wide range of primary cell types and cell lines^11;39^. We also included as annotations the output from “genome segmentation” of the six main ENCODE cell lines^40^; for each section of the genome in each cell line, Hoffman et al.^40^ report whether the histone modifications in the region are consistent with enhancer activity, transcription start sites, promoter-flanking regions, CTCF binding sites, or repressed chromatin. Finally, we included elements of gene structures (protein-coding exons and 3’ and 5’ UTRs). In total, we used data from 18 traits and 450 genomic annotations.

For each trait, we set out to identify which of the 450 annotations (if any) were enriched in genetic variants influencing the trait. To do this, we developed a hierarchical model that learns the shared properties of loci influencing a trait. The full details of the model are presented in the Methods, but can be summarized briefly. Conceptually, we break the genome into large, non-overlapping blocks (with an average size of 2.5 Mb). Let the prior probability that any block *k* contains an association be Π*_k_*. *If* there is an association in block *k*, then let the prior probability that any SNP *i* is the causal SNP be *π_ik_*. We allow both Π*_k_* and *π_ik_* to depend on annotations of the region and SNP, respectively, and estimate these quantities based on the patterns of enrichment across the whole genome. We tested this approach using simulations based on real data from a GWAS of height (Supplementary Material).

The methodology is best illustrated with an example. We started with an analysis of a GWAS of high-density lipoprotein (HDL) levels^27^. We first took each genomic annotation individually, and estimated its level of enrichment (or depletion) for loci that influence HDL (in the model, we additionally included a regional effect of gene density and and a SNP-level effect of distance to the nearest transcription start site, see the Methods for details). In Figure 1A, we show the top 40 annotations, ordered by how well each improves the fit of the model. Loci that influence HDL are most strongly enriched in enhancers identified in the HepG2 cell line, and most strongly depleted from genomic regions repressed in that same cell line. HepG2 cells are derived from a liver cancer; the relevance of this cell line to a lipid phenotype makes intuitive sense. However, there are many other additional (correlated) genome annotations that are enriched for loci that influence HDL (Figure 1A). We thus built a model including multiple annotations; to mitigate over-fitting in this situation we used a cross-validation approach (Methods). The best-fitting model is shown in Figure 1B. It include both enhancers and repressed chromatin identified in HepG2 cells, as well as coding exons and chromatin repressed in K562 cells. Since many of the annotations are correlated, those included in the combined model are the “best” representatives of sets of related annotations. We thus used conditional analysis to define the set of annotations represented by each member of the combined model (Methods; Figure 1A).

**Figure 1.**
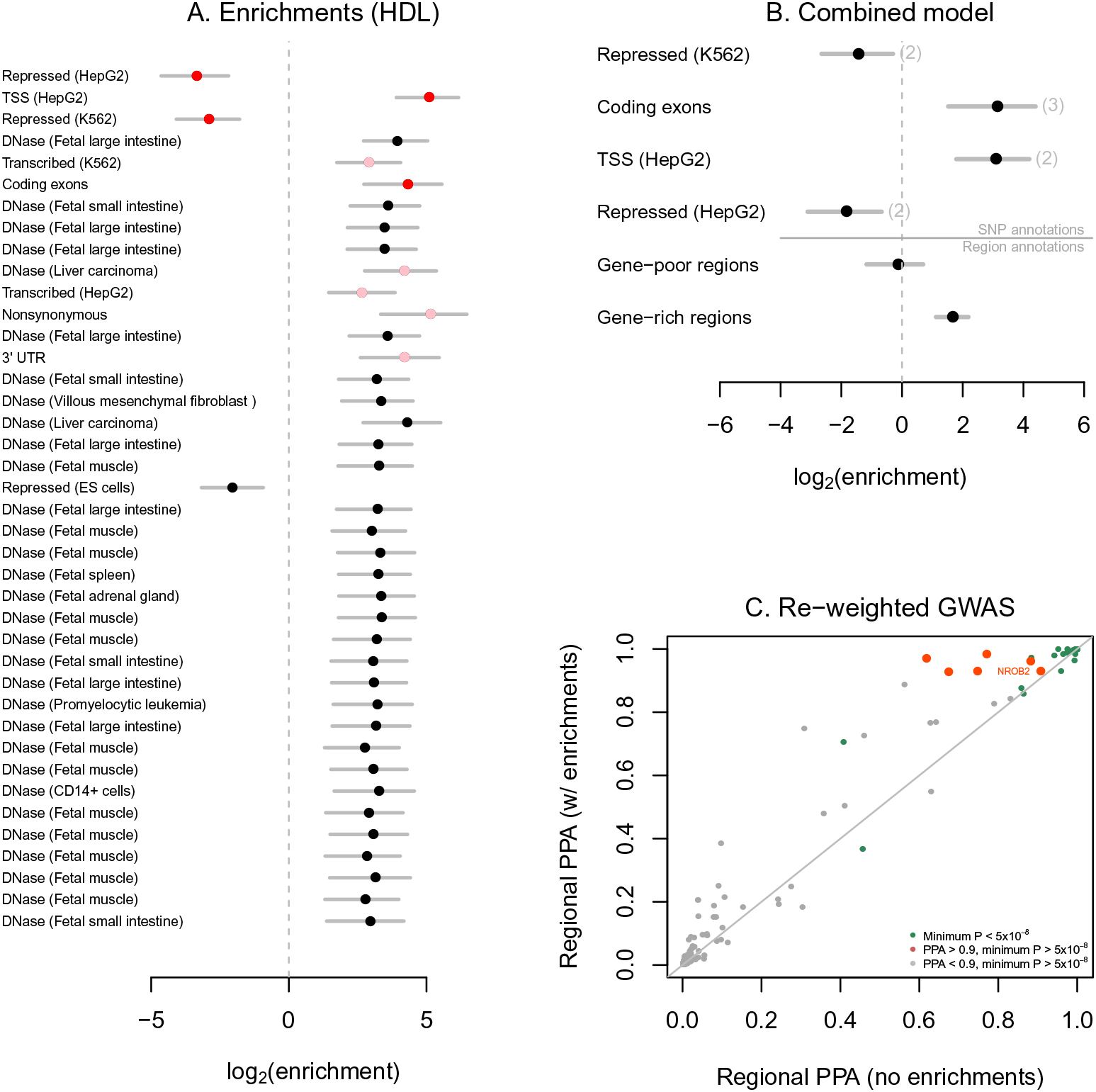
Application of the model to HDL cholesterol. A. Single-annotation models. We fit the model to each annotation individually, including a SNP-level effect for SNPs 0-5 kb from a TSS, a SNP-level effect for SNPs 5-10 kb from a TSS, a region-level effect for regions in the top third of gene density, and a region-level effect for regions in the bottom third of gene density. Plotted are the maximum likelihood estimates and 95% confidence intervals of the enrichment parameter for each annotation. Annotations are ordered according to how much they improve the likelihood of the model (at the top are those that improve the likelihood the most). In red are the annotations included in the joint model, and in pink are the annotations that are statistically equivalent to those included in the combined model. **B. Joint model.** Using the algorithm described in the Methods we built a model combining multiple annotations. Shown are the maximum likelihood estimates and 95% confidence intervals of the enrichment effects of each annotation. Note that though these are the maximum likelihood estimates, model choice was performed using a penalized likelihood. In parentheses next to each annotation (expect for those relating to distance to transcription start sites), we show the total number of annotations that are statistically equivalent to the included annotation in a conditional analysis. **C. Re-weighted GWAS.** We re-weighted the GWAS using the model with all the annotations in **B** (under the penalized enrichment parameters from Supplementary Table 9). Each point represents a region of the genome, and shown are the posterior probabilities of association (PPA) of the region in the models with and without the annotations.

A convenient side effect of fitting an explicit statistical model relating properties of SNPs to the probability of association is that we can use the functional information to re-weight the GWAS (Methods). We used the combined model for HDL to re-weight the association statistics across the genome (Figure 1C). There are several regions of the genome with strong evidence for association with HDL (posterior probability of association [PPA] over 0.9) only when using the model incorporating functional information. In Figure 2, we show one such region, near the gene NR0B2. The model identifies the SNP rs6659176 as the most likely candidate to be the causal polymorphism in this region. This SNP has a P-value of 1.5 × 10^−6^. However, this SNP falls falls in a coding exon (in fact it is nonsynonymous), leading the model to conclude that this P-value is in fact strong evidence for association. Indeed, larger studies of HDL have confirmed the evidence for association in this region (P-value of 9.7 × 10^−16^ at rs12748152, which has *r*^2^ = 0.85 with rs6659176^17^). This region, though not this particular SNP, was also identified in a scan for SNPs influencing multiple lipid phenotypes^41^.

**Figure 2.**
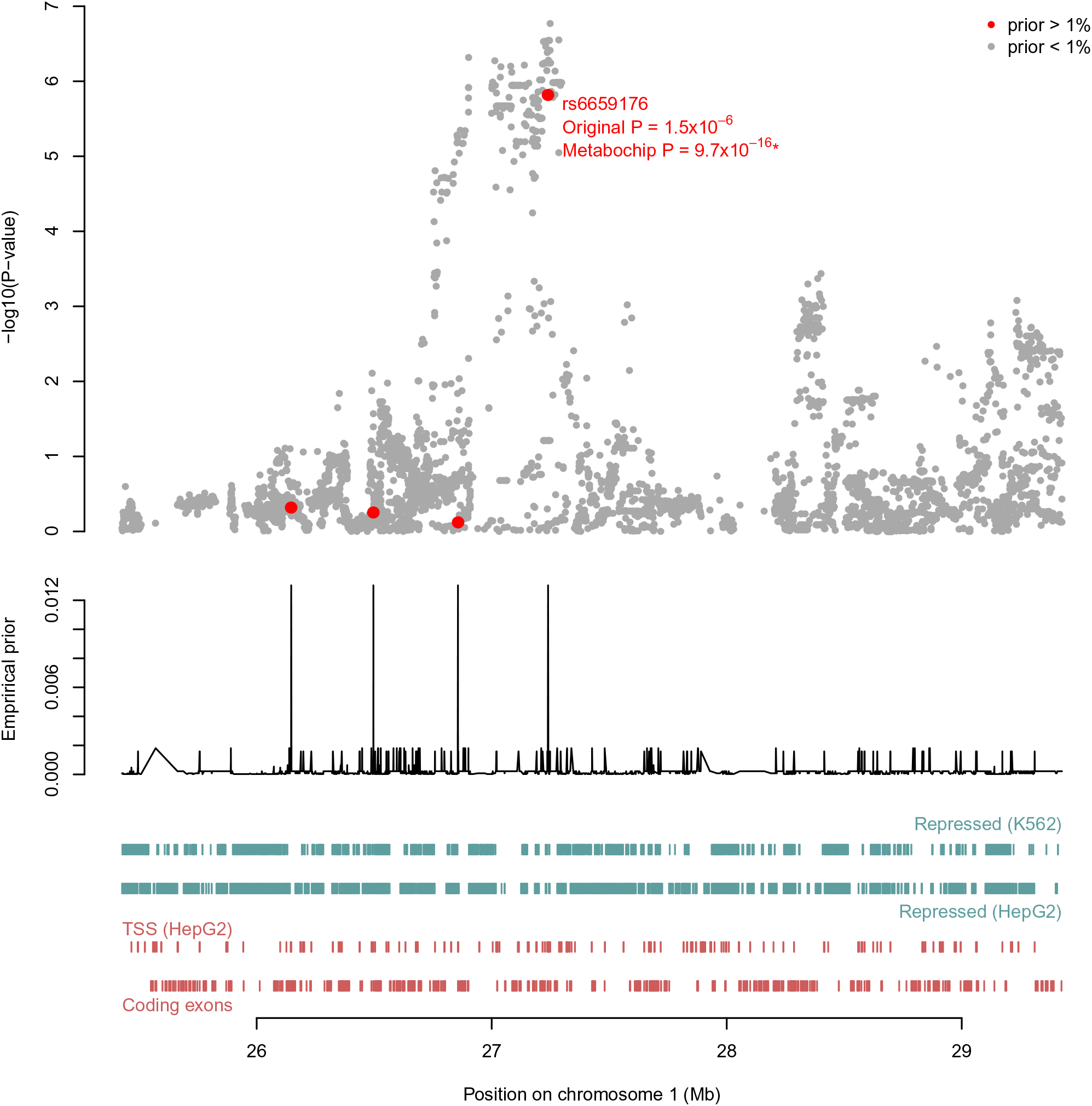
Regional plot surrounding NR0B2. In the top panel we plot the P-values for association with HDL levels at each SNP in this region. In the middle panel is the fitted empirical prior probability that each SNP is the causal one in the region, conditional on there being a single causal SNP in the region. This prior was estimated using the combined model with the annotations in Figure 1B. In the lower panel are the positions of the annotations included in the model.* The reported P-value is for rs12748152, which has *r*^2^ = 0.85 with rs6659176.

We applied this method to all 18 traits. We were first interested in estimating the fraction of associations for each trait that can be explained by nonsynonymous polymorphisms versus polymorphisms that do not influence protein sequences. For each trait, we fit a model including promoters (SNPs within 5 kb of a transcription start site) as well as nonsynonymous polymorphisms. For all traits, nonsynonymous polymorphsims are enriched among those that influence the trait, though this enrichment is not statistically significant for all traits (Figure 3A). This contrasts with synonymous polymorphisms, which are generally not enriched for polymorphisms that influence traits, with a few notable exceptions (like height and Crohn’s disease, see Supplementary Figure 3). We then used these enrichments to estimate the fraction of associations for each trait that are driven by nonsynonymous polymorphisms (Supplementary Material). This fraction varies from around 2% to around 20%, with an average of 10% (Figure 3B). We conclude that the relative importance of changes in protein sequence versus gene expression likely varies across traits.

**Figure 3.**
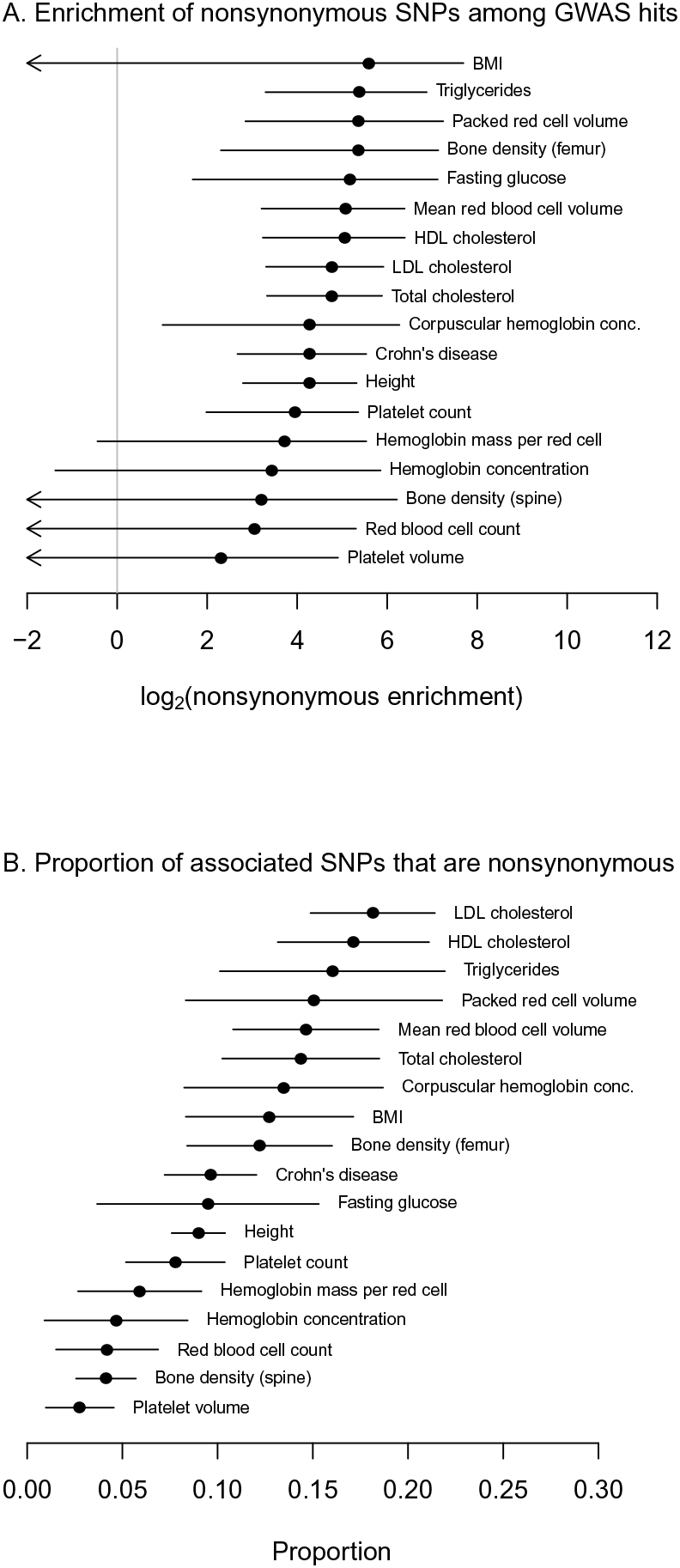
Estimated role of protein-coding changes in each trait. **A. Estimated enrichment of non-synonymous SNPs.** For each trait, we fit a model including an effect of non-synonymous SNPs and an effect of SNPs within 5 kb of a TSS. Shown are the estimated enrichments parameters and 95% confidence intervals for the non-synonymous SNPs. **B. Estimated proportion of GWAS hits driven by non-synonymous SNPs.** For each trait, using the model fit in **A.**, we estimated the proportion of GWAS signals driven by non-synonymous SNPs. Shown is this estimate and its standard error (Supplementary Material).

We then used all 450 genome annotations to build models of enrichment for each trait. As for HDL, we first estimated enrichment levels individually for each annotation (Supplementary Figures 4-12). Clustering of phenotypes according to these enrichment levels recapitulated many known relationships between traits (Supplementary Figure 13). We then generated a combined model for each trait. The parameters of the combined models are shown in Figure 4 and Supplementary Figure 14, and details of the exact annotations are in Supplementary Tables 2-20. In general, the models generated with this method are sparse and biologically interpretable. A few general patterns emerge from this analysis. Apart from the repeated occurrence of annotations related to protein-coding genes, marks of repressed chromatin are often significantly depleted for SNPs influencing traits. For example, SNPs influencing Crohn’s disease are depleted from repressed chromatin identified in a lymphoblastoid cell line (Figure 4D; log_2_ enrichment of -1.83, 95% CI [-3.06, -0.78]), SNPs influencing height are significantly depleted from repressed chromatin in HeLa cells (Figure 4I; log_2_ enrichment of -1.5, 95% CI [-2.39, -0.71]), and SNPs influencing red blood cell volume are significantly depleted from repressed chromatin in an erythroblast-derived cell line (Figure 4F; log_2_ enrichment of -3.91, 95% CI [-6.25, -2.38]).

**Figure 4.**
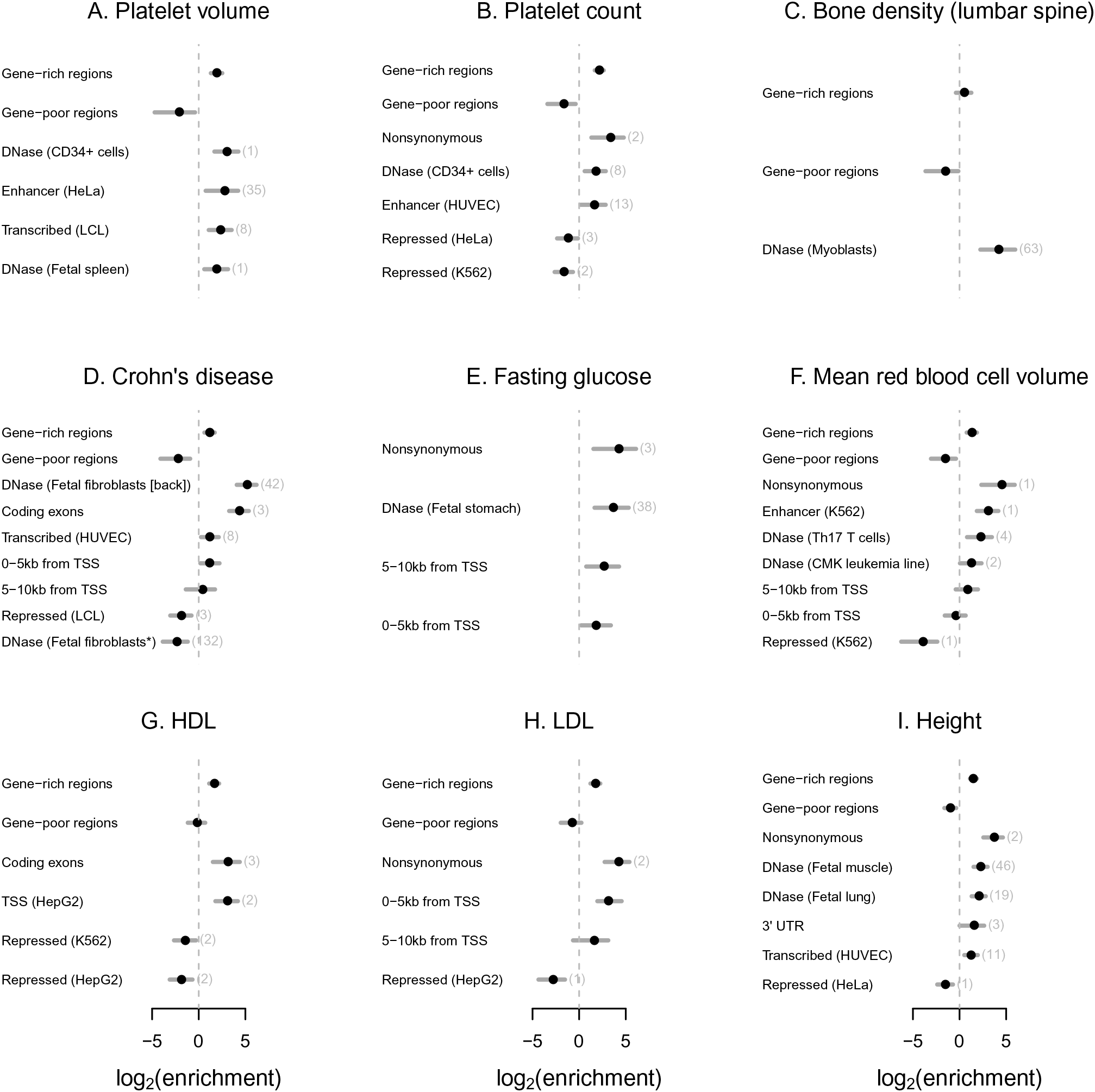
Combined models for nine traits. For each trait, we built a combined model of annotations using the algorithm presented in the Methods. Shown are the maximum likelihood estimates and 95% confidence intervals for all annotations included in each model. Note that though these are the maximum likelihood estimates, model choice was done using a penalized likelihood (Methods). For the other nine traits, see Supplementary Figure 14. In parentheses next to each annotation (expect for those relating to distance to transcription start sites), we show the total number of annotations that are statistically equivalent to the included annotation in a conditional analysis (Methods). *This annotation of DNase-I hypersensitive sites in fetal fibroblasts from the abdomen has a positive effect when treated alone; see the text for discussion.

We additionally observed cell type-specific enrichments in enhancer elements and DNase hypersensitive sites for SNPs that influence traits. Most of the observed enrichments are readily interpreted in light of the known biology of the trait. For example, SNPs that influence platelet volume and platelet count are enriched in open chromatin identified in CD34^+^ cells, known to be on the cell lineage that leads to platelets^42^ (Figure 4A,B; log_2_ enrichment of 1.81, 95% CI [0.59, 2.86] for platelet count; log_2_ enrichment of 3.02, 95% CI [1.69, 4.26] for platelet volume); and SNPs that influence corpuscular hemoglobin concentration are enriched in open chromatin identified in K562 cells, a cell line derived from a cancer of erythroblasts (Supplementary Figure 14E; log_2_ enrichment of 2.67, 95% CI [0.61, 4.44]). For some traits, however, the connection between the trait and the tissues identified is not immediately obvious. For example, SNPs associated with platelet density are enriched in open chromatin in the spleen (Figure 4A; log_2_ enrichment of 1.93, 95% CI [0.59, 3.14]), and SNPs associated with height are enrichment in open chromatin in muscle (Figure 4I; log_2_ enrichment of 2.27, 95% CI [1.51, 3.02]; note that though there are all a large number of annotations equivalent to this annotation of open chromatin in fetal muscle, all of them are muscle-related; see Supplementary Figure 5B).

For two traits–Crohn’s disease (Figure 4D) and red blood cell count (Supplementary Figure 14G)–we noticed that annotations initially identified as enriched for SNPs influencing the trait ended up in the combined model as being depleted for SNPs influencing the trait. On further examination (Supplementary Material), we found that these effects are due to statistical interactions. For example, when treated alone, SNPs that influence Crohn’s disease are enriched in DNase-I hypersensitive sites identified in fetal fibroblasts from the abdomen (log_2_ enrichment of 3.17, 95% CI [1.66, 4.36]). However, DNase-I hypersensitive sites identified in fetal fibroblasts from the back show an even stronger enrichment (log_2_ enrichment of 4.21, 95% CI [2.99, 5.29]), and sites in common between the two annotation are intermediate (log_2_ enrichment of 3.74, 95% CI [2.29, 4.89]). This leads to an interaction where in the joint model the contribution of the DNase-I hypersensitive sites identified in fetal abdominal fibroblasts is negative. Though this is a statistical explanation for this observation, the biological explanation is not immediately clear. It seems likely that DNase-I hypersensitive sites are a heterogeneous set of different classes of elements, and that different experiments are more sensitive, for either technical or biological reasons, to subsets of these elements.

Finally, we explored the potential of this model to identify new loci (as in Figure 2). In order to do this, one needs a threshold for “significance” in this model, ideally with similar properties as the standard P-value threshold of 5 × 10^−8^. To calibrate the method, we used the fact that we initially applied our method to a study of four lipid traits that identified about 100 loci in a sample size of around 90,000 individuals^27^. Since then, larger studies have raised the number of loci associated with lipid traits to 157^17^. If we treat a locus with a P-value of 5 × 10^−8^ in the larger study as a “true positive” and a locus that does not reach this threshold in the larger study as a “true negative”, we can calibrate a threshold for posterior probability of association using the replication data (Supplementary Material). We found that a threshold of a regional PPA of 0.9 performed similarly to a stringent P-value threshold (Supplementary Figure 15). Combining the loci from both the standard P-value approach and our approach resulted in an approximately 5% increase in the number of identified loci while still maintaining a false positive rate close to zero (Supplementary Figure 15, Supplementary Table 21). This is only a modest gain in power; that said, by applying this method to all 18 traits we identified 49 loci that did not reach a standard statistical significance threshold of 5 × 10^−8^ but have a PPA over 0.9 (Supplementary Table 22). Based on the above results for lipids, the level of evidence that these loci are true positives is approximately the same as those that have *P* = 5 × 10^−8^ in a standard GWAS. Indeed, the majority of these loci have since been identified in larger cohorts than those used in this paper (Supplementary Table 22).

## 3 Discussion

In this paper, we have developed a statistical model for identifying genomic annotations that are most relevant to the biology of a given phenotype. We have shown that this model is able to scan through hundreds of genomic annotations to identify a sparse set of biologically-interpretable annotations without prior knowledge of the biology of the phenotype.

### Linking GWAS to biology

Perhaps the most striking observation is that chromatin annotated as repressed in a given cell type is often depleted for SNPs that influence traits. Since approximately 60-70% of the genome falls in this annotation in any given cell type (Supplementary Material), this information could dramatically limit the number of SNPs considered when fine-mapping loci identified in GWAS. Additionally, we identified several non-obvious connections between tissue and phenotypes. For example, SNPs that influence platelet volume are enriched in DNAse-I hypersensitive sites in the spleen (Figure 4A). Though the spleen functions in the removal of platelets from the bloodstream, the connection between its function and platelet volume is unclear. An important next step will be to connect the identified non-coding variants in regulatory regions to changes in gene expression, which is presumably the mechanism by which they exert an effect on phenotypes. Methods for inferring the casual chain between variation in DNAse-I sensitivity, variation in gene expression, and variation in phenotypes (e.g.^41;43–46^) will be essential.

### Modeling assumptions

We have made several modeling assumptions that merit discussion. First, by splitting the genome into blocks based on numbers of SNPs, we are making the implicit assumption that the probability that a genomic region contains a SNP associated with a given phenotype depends on the SNP density rather than the physical size–that is, a short genomic region with a large number of SNPs is *a priori* as likely to have an association as a long genomic region with few SNPs. We have also made a more restrictive assumption that there can be only a single causal SNP in a given genomic region. This assumption is a natural starting point, but as GWAS sample sizes increase even more it will begin to be untenable. Advances in methods for joint analysis of multiple SNPs (e.g. Yang et al.^47^) may provide a way forward in this situation. Finally, we note that the model is limited by the types of genomic annotations that are available, and the best annotations identified in the model may be “proxies” for the truly-relevant annotations. For example, SNPs associated with height are enriched in DNase-I hypersensitive sites identified in the fetal lung (Figure 4I); taken literally, this would suggest that some SNPs influence height though lung development. An alternative possibility, however, is that patterns of open chromatin in the lung (which is of course a heterogeneous tissue) are useful proxies for patterns of open chromatin in a cell type that has not been profiled; this hypothetical cell type could in principle be present in any tissue.

### Prospects for fine-mapping GWAS loci using functional genomic data

We have primarily focused on using our model to identify annotations relevant to a trait of interest, though we have also explored using this information to identify novel loci. A third natural application, which we have not explored, is the possibility for fine-mapping GWAS loci using functional genomic information^48^. Indeed, the posterior probability that each SNP in a given genomic region is the causal one is explicitly included in our model. However, in current applications around 20% of common SNPs are neither genotyped nor successfully imputed; this is a major limitation to fine-mapping that cannot be overcome with statistical means. As GWAS move to even denser genotyping or sequencing, we expect that re-visiting this issue will be fruitful.

## Methods

In this section we detail the specifics of the hierarchical model; a description of the data used is in the Supplementary Material. The model we propose is most closely related to that developed by Veyrieras et al.^23^ in the context of eQTL mapping. Conceptually, we split the genome into independent blocks, such that the blocks are larger than the extent of LD in the population. We then allow each block to contain either a single polymorphism that causally influences the trait or none. We model the prior probability that any given block contains an association and the conditional prior probability that any given SNP in the block is the causal one. The key is that we allow these probabilities to vary according to functional annotations–for example, gene-rich regions might be more likely to contain associations, and if there is an association, the causal polymorphism may be more likely to fall in a transcription factor binding site. We then estimate these priors using an empirical Bayes approach. Software implementing this model is available at https://github.com/joepickrell/fgwas.

### Computing the Bayes factor

The basic building block of the model is a linear regression model. Consider a single SNP genotyped in *N* phenotyped individuals. Assume each individual has an associated measurement of a quantitative trait (we describe a slight modification for case-control studies later), and let 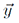 be the vector of phenotypes. Let 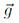 be the vector of genotypes (coded 0, 1, or 2 according to counts of an arbitrarily-defined allele). We use a standard additive linear model:

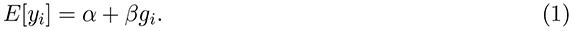

We would like to compare two models: one where *β* = 0 and one where *β* ≠ 0. A natural way to compare these two models is the Bayes factor:

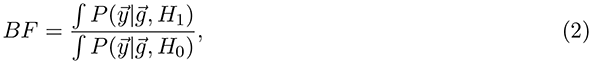

where *H*_1_ and *H*_0_ represent the parameters of the the alternative and null models, respectively, and which are integrated out.

To compute Equation 2, we use the approximate Bayes factor from Wakefield^49^. This Bayes factor has the practically important property that is can be calculated from a summary of the linear regression, without access to the underlying genotype vector 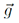. For completeness, we re-iterate here the underlying model. If 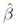 is the maximum likelihood estimator of *β* and 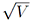 is the standard error of 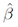, Wakefield^49^ suggests a model in which:

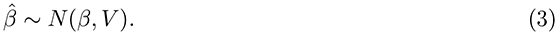

Wakefield^49^ places a normal prior on *β*, such that *β ∼ N* (0*, W)*. Under this model Equation 2 becomes:

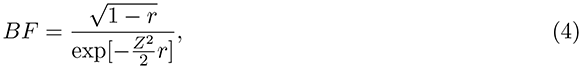

where 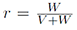 and 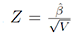 (a standard Z-score). Thus, from a Z-score, an estimate of *V*, and the prior variance *W*, we can obtain a Bayes factor measuring the statistical support for a model in which a SNP is associated with a trait as compared to a model in which a SNP is not associated with a trait. Note, however, that because of linkage disequilibrium in the genome, any true causal association will lead to multiple true statistical associations. In all applications, we set *W* = 0.1 as the prior, such that the majority of the weight of the prior is on small effect sizes (results are robust to some variation in this prior; see Supplementary Material and Supplementary Figure 16).

### Hierarchical model

Now consider a set of *M* SNPs, each of which has been genotyped in *N* individuals in a GWAS. Our goal is to build a model to identify the shared characteristics of SNPs that causally influence a trait. Because of LD, there will be many associations in the genome that are not causal; however, these will all be restricted to a block around the truly causal site. We thus split the genome into contiguous blocks of size *K* SNPs (in all of our applications, we set *K* = 5, 000, though doubling this block size had little effect on the results; see Supplementary Material and Supplementary Figure 16), such that there are *M/K* blocks. We choose the block size to be much larger than the extent of linkage disequilibrium in the population. Let Π*_k_* be the prior probability that block *k* contains a causal SNP associated with the trait. The probability of the data (the set of observed phenotypes) is then:

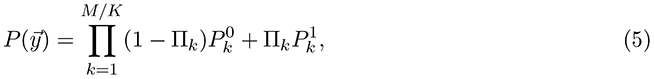

where 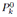 is the probability of the data in block *k* under the model where there are no SNPs associated with the trait in the block, and 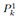 is the probability of the data in block *k* under the model where there is one SNP associated with the trait in the block. Further,

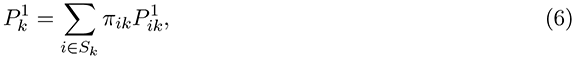

where *S_k_* is the set of SNPs in block *k*, *π_ik_* is the prior probability that SNP *i* is the causal SNP in the region conditional on there being an association in block *k*, and 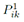 is the probability of the data under the model where this SNP is associated with the trait. Note that this is not a multiple regression model where we jointly model the effects of multiple SNPs on a trait (as in, for example, Carbonetto and Stephens^31^).

We can now allow the prior probabilities–both Π*_k_* (the prior on the block of SNPs containing an association) and *π_ik_* (the prior probability that SNP *i* is the causal SNP assuming there is a single association in block *k*)–to depend on external information. We would also like to avoid subjective variation in Π*_k_* and *π_i_*, but instead learn from the data itself which genomic annotations are most important. Specifically, we model the regional prior probability as:

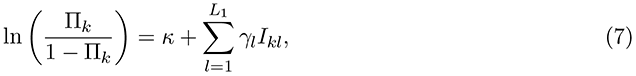

where *L*_1_ is the number of region-level annotations in the model, *γ_l_* is the effect associated with annotation *l* and *I_kl_* takes the value 1 if region *k* is annotated with annotation *l* and 0 otherwise. For example, in practice we will estimate a *γ* parameter for regions of high or low gene density. We then model the SNP prior probability as:

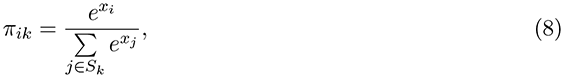

where

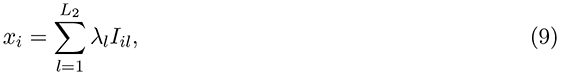

where *L*_2_ is the number of SNP-level annotations in the model, *λ_l_* is the effect of SNP annotation *l* and *I_il_* takes the value 1 if SNP *i* falls in annotation *l* and 0 otherwise. For example, in practice we will estimate a *λ* parameter for nonsynonymous SNPs.

### Fitting the model

Combining terms above, we see that the likelihood of the data can be written down as:

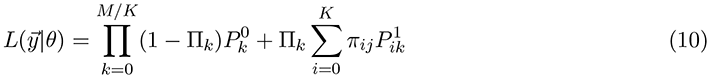

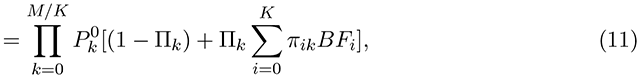

where *θ* is contains all the parameters of the model, most notably the set of annotation parameters. We maximize this function using the Nelder-Mead algorithm implemented in the GNU Scientific Library.

### Shrinkage estimators of the annotation parameters

While maximizing Equation 10 gives the maximum likelihood estimates of all parameters, one concern is that there may be some level of overfitting. When comparing models, we instead shrink these parameters towards zero. Specifically, we define a penalized log-likelihood function:

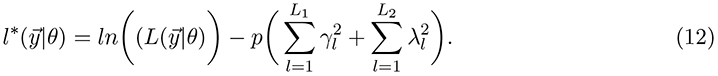

The penalty *p* on the sum of the squared annotation parameters is the one used in ridge regression^50^. In ridge regression, parameter estimates under this penalty are equivalent to estimating the posterior mean of the parameter if the prior distribution of the parameter is Gaussian^50^; changing the tuning parameter *p* is equivalent to changing the prior. We suspect that the interpretation in this model is similar. Since this penalized likelihood can not be used for formal statistical tests, we tune the *p* parameter by cross-validation. An alternative approach here would be to explicitly put a prior on the enrichment parameters, but in the absence of a conjugate prior this would likely add substantially to the computational burden for little practical benefit.

### Cross-validation

To compare models and tune the penalty *p* in the penalized likelihood above, we used a 10-fold cross-validation approach. We split the chromosomal segments into 10 folds. Let 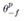 be the parameters of the model estimated while holding out the data from fold *f* and under penalty *p*, and 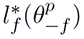 be the penalized log-likelihood of the data in fold *f* under the model optimized without using fold *f*. Then:

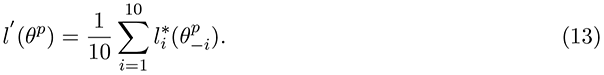

Note that the size of the folds used in this cross-validation means that each fold excludes more than an entire chromosome. This means that no individual chromosome can have undue influence on the parameters included in the model.

### Model choice

Consider a single phenotype and a set of *L* functional annotations of SNPs (in our case *L* is in the hundreds). Including all *L* SNP annotations in the model is neither biologically interesting nor computationally feasible. We thus set out to choose a relatively sparse model that fits the data. We start with forward selection: for each of the *L* annotations, we fit a model including a region-level parameter for regions in the top third of the distribution of gene density, a region-level parameter for regions in the bottom third of the distribution of gene density, a SNP-level parameter for SNPs from 0-5 kb from a TSS, a SNP-level parameter for SNPs from 5-10 kb from a TSS, and a SNP-level parameter for the annotation in question. We then identify the set of annotations that significantly improve the model fit (as judged by the likelihood from Equation 10). We then:

1. Add the annotation that most significantly improves the likelihood to the model.
2. For each annotation identified as having a significant marginal effect, test a model including the annotation and those that have already been added.
3. If any annotation remains significant, go back to step 1.

At this point, there are generally a small number of annotations in the model, but the model may be over-fit. We then switch to using the 10-fold cross-validation likelihood in Equation 13. We first tune the penalty parameter *p* by finding the value of *p* that maximizes the cross-validation likelihood. We then:

1. Drop each annotation from the model in turn, and evaluate the cross-validation likelihood. When dropping annotations, we additionally try dropping the region-level annotations on gene density and the SNP-level annotations on distance to the nearest TSS.
2. If a simpler model has a higher cross-validation likelihood than the full model, drop the annotation from the model and return to step 1.
3. Report the model that has the highest cross-validation likelihood.

### Approximating ***V_i_***

In order to compute the Bayes factor in Equation 4, we need an estimate of *V_i_*, the standard error of the estimated effect size of SNP *i*. In principle, this is trivial output from standard regression software; however, it is rarely reported. Instead, let *f_i_* be the minor allele frequency of SNP *i* computed from an external sample of the same ancestry as the population in which the association study was done (we use data from the 1000 Genomes Project^38^). Let *N_i_* be the number of individuals in the association study at SNP *i* (this can vary across SNPs due to missing data). Then:

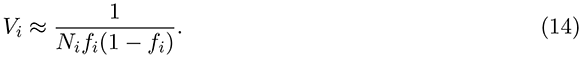

Note that this variance is independent of the actual scale of the measurements; this is appropriate because the Z-scores are independent of the scale of the measurements as well.

### Case-control studies

For all of the above, we have considered studies of quantitative traits. For a case-control study, we assume that we have summary statistics from logistic regression instead of linear regression. All aspects of the model are identical, with the exception of the approximation of *V_i_*. Define *N_case_* and *N_control_* as the numbers of cases and controls, respectively. Now, ^49^:

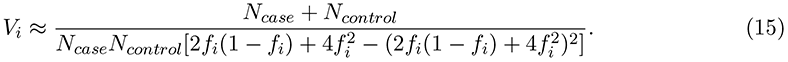

The variance here is on a log-odds scale.

### Posterior probabilities of association

Once the model has been fit, we have empirical estimates of the prior probability that region *k* contains an association, 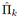 and the prior probability that SNP *i* is the causal one, 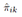 (conditional on there being an association). We define a Bayes factor summarizing the evidence for association in the *region* (see, for example Maller et al.^48^):

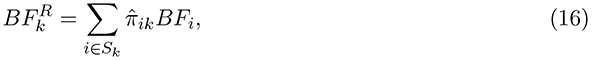

where *S_k_* is the set of SNPs in region *k* and *BF_i_* is the Bayes factor for SNP *i* (Equation 4). The posterior probability that region *k* contains an association is then:

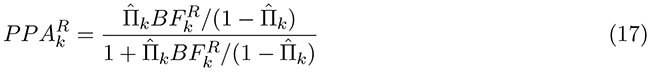

We can also define the posterior probability that any given SNP *i* in region *k* is the causal one under our model:

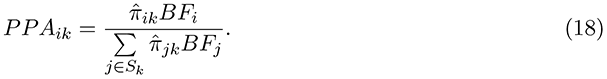

This is similar to the calculation in Maller et al.^48^, except that we allow the prior probability *π_ik_* to vary across SNPs.

Finally, we can define the posterior probability that any given SNP is causal. This is the posterior probability that the region contains a causal SNP times the posterior probability that the SNP is causal conditional on there being an association in the region. If SNP *i* falls in region *k*:

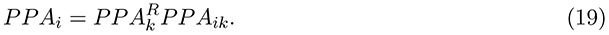

### Conditional analysis

Because many of the annotations we consider are correlated, those ultimately included in the combined model for each trait (Figure 4) may be representatives of a large group of correlated annotations. For biological interpretation of the model, it is thus important to know which of the other annotations are interchangeable with those included in the model.

To test this, we took an approach of conditional analysis. Consider two SNP-level annotations, with annotation parameters *λ*_1_ and *λ*_2_, respectively. In a joint model, we would jointly estimate both *λ*_1_ and *λ*_2_. However, we are interested in whether the second annotation adds information above and beyond that provided by the first annotation. We thus first estimate *λ*_1_, then *fix* this parameter to its maximum likelihood value 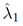. We then estimate 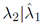 that is, we obtain the maximum likelihood estimate and 95% confidence interval of *λ*_2_ *conditional* on a fixed value of *λ*_1_. If this confidence interval does not overlap zero, this is evidence that the second annotation adds information to the model above that provided by the first.

In practice, we first fit the combined model as described in the section “Model choice” above. We then returned to the set of annotations that had significant marginal associations. For each annotation in the combined model, we took each of the other annotations in turn and tested whether the included annotation was significantly more informative than the non-included annotation. In Figure 4 and Supplementary Figure 12, we display the total number of annotations represented by each one that is included in the combined model.

## Acknowledgements

We thank David Reich, Nick Patterson, Alkes Price, and Po-Ru Loh for helpful discussions and suggestions; and Jonathan Pritchard, Graham Coop, and two anonymous reviewers for comments on a previous version of this manuscript. We thank Nicole Soranzo for providing access to the platelet studies, and Luke Jostins for assistance in obtaining the Crohn’s disease data. This work was supported by NIH postdoctoral fellowship GM103098 to JKP.

